# Released mitochondrial DNA and neurofilament light chain as Parkinson’s disease phenotypes in patient-specific midbrain assembloids

**DOI:** 10.1101/2025.03.28.645921

**Authors:** Sonia Sabate-Soler, Isabel Rosety, Gemma Gomez-Giro, Jenny Ghelfi, Soraya Hezzaz, Anne Grünewald, Jens C. Schwamborn, Javier Jarazo

## Abstract

Parkinson’s Disease is the second most common neurodegenerative disorder worldwide, with growing numbers and considerable societal and economic concerns. Human cell culture systems are efficient models for neurodegenerative disorders and allow for personalized, non-invasive analysis of cellular and molecular disease mechanisms. Midbrain organoids and assembloids are advanced 3D culture systems that recapitulate the human midbrain, which is highly affected by Parkinson’s disease. Here, we used healthy control and patient-specific midbrain assembloids to assess mitochondrial DNA phenotypes and NfL levels alongside neurodegeneration and alpha-synuclein phosphorylation. Importantly, alterations in mitochondrial DNA homeostasis and NfL levels can be assayed in the supernatant and therefore are particularly suitable as biomarkers and for high throughput screening approaches.

## Introduction

Parkinson’s Disease (PD) is the second most common neurodegenerative disorder worldwide, affecting around 3% of the worldwide population over the age of 80 [1]. Different factors play a role in the development of PD, including genetics – such as mutations in the *SNCA, LRRK2*, and *PINK1* genes – and environmental factors – such as pesticide exposure. Monogenic forms of PD account for 20% of the cases, while 80% are idiopathic [2].

Mitochondrial dysfunction is one of the pathological processes described in PD. Impaired mitochondrial function and metabolism have been observed in idiopathic and monogenic PD forms. Mutations in genes such as *PINK1* and *DJ-1* have been linked to mitochondrial dysfunction in PD [3], [4].

One of the main pathophysiological hallmarks of PD is the accumulation of aggregated alpha-synuclein in dopaminergic neurons. The relationship between alpha-synuclein and mitochondrial homeostasis has been studied, and proven evidence of their relation and functionality has been shown. [5], [6]. Mitochondrial homeostasis is not only related to alpha-synuclein aggregation but also to neurodegeneration. Studies conducted over the last few years point to a direct relationship between mitochondrial dysfunction, neuroinflammation, and neurodegeneration. [7], [8]

Midbrain organoids (MOs) are advanced, iPSC-derived, 3D cell culture systems that mimic the human midbrain [9]. Co-culturing MOs with iPSC-derived macrophage precursors leads to microglia-containing assembloids, which have a heterogeneous cellular composition and show neuronal functionality [10]. MOs recapitulate PD phenotypes and can be used for drug testing [11].

In this article, we use microglia-containing assembloids as a PD model. We observed phenotypes related to neurodegeneration, as well as alpha-synuclein phosphorylation. We analyzed cell culture supernatants from assembloids with a wild-type (WT) background or from individuals carrying a triplication in the *SNCA* gene (3x SNCA), observing neurodegeneration and mitochondrial homeostasis phenotypes. Importantly, a significant release of Neurofilament light chain (NfL) was detected in midbrain assembloids. Furthermore, mitochondrial DNA copy number and transcription alterations were detected in those supernatants, opening the door to biomarker detection in cell culture media from human-personalized organoid and assembloid models.

## Material and Methods

- Assembloid generation Human midbrain organoids were generated as described in [12], [13]. In short, 9000 neuro-epithelial stem cells (NESCs) were seeded per well in ultra-low attachment 96-well plates (FaCellitate, F202003). They were cultured for 2 days in maintenance medium (N2B27 medium supplemented with 150 μM Ascorbic acid [Sigma, A4544], 3 μM CHIR 99021 [Axon Medchem, 1386], 3 μM Purmorphamine [PMA, Enzo Lifesciences, ALX-420-045-M001]). After that, the medium was switched to neuronal differentiation-inducing medium (N2B27 medium supplemented with 200 μM Ascorbic acid [Sigma, A4544], 10 ng/ml Brain Derived Neurotrophic Factor [BDNF, Peprotech, 450-02], 10 ng/ml Glial-Derived Neurotrophic Factor [GDNF, Peprotech, 17814463], 1 ng/ml TGF-β3 (Peprotech, 100-36E), 500 μM db cAMP [Biosynth, D07996], 1 μM Purmorphamine [PMA, Enzo Lifesciences, ALX-420-045-M001]). The organoids were cultured with neuronal differentiation-inducing medium for 5 days. The day after, organoids were co-cultured with macrophage precursor cells following [10], with some modifications, in order to generate assembloids. Shortly, 50,000 macrophage precursor cells were added to each developing organoid, in co-culture medium (Advanced DMEM/F12 [Gibco, 12634028], 1x N2 supplement [Thermofisher, 17502001], 1x GlutaMAX™ [Thermofisher, 35050061], 50 μM 2-mercaptoethanol [Thermo Scientific, 11528926], 100 U/ml Penicillin– Streptomycin [Gibco, 15140163], 100 ng/ml IL-34 [Peprotech, 200-34], 10 ng/ml GM-CSF [Peprotech, 200-34], 10 ng/ml BDNF, 10 ng/ml GDNF), and assembloids were cultured with co-culture medium for an additional 53 days.
- Media collection and storage Assembloid culture media was collected in 96 Well Polypropylene Sample Storage Microplates (Thermo Scientific, 267334), and sealed with aluminum plate seals (Thermo Scientific, 232698). After that, the collected samples were snap-frozen and stored at -80C before analysis.
- Mitochondrial DNA analysis
  ∘ DNA extraction The assembloid medium was collected and snap-frozen for storage. 90 µL of culture medium was collected per assembloid, and before extraction, samples were diluted by adding 110 µL of 1x PBS (Gibco, 14040-091). 20 µL Proteinase K (Qiagen, RP103B) at a concentration of 40mAU/mg or 600mAU/mL was added to each sample, and samples were processed following manufacturer instructions for the QIAamp DNA Mini Kit (Qiagen, 51304).
  ∘ Digital PCR Digital PCR (dPCR) was performed using QuantStudio™ Absolute Q™ Digital PCR System (Applied Biosystems, A52864). The primers used for mitochondrial *ND1, ND4*, D-loop and *B2M* are described in [14]. Samples were prepared by adding 2 µL of the 5x Absolute Q™ DNA Digital PCR Master Mix (Applied Biosystems, A52490), 0,166 µL of the corresponding 60x primer probes, and MilliQ water to 5 µL of DNA, in a final volume of 10 µL per reaction. The triplex qPCR program was run, with an initial 10-minute run at 96°C followed by 40 cycles of 5 seconds at 96°C and 30 seconds at 60°C.
- NfL ELISA Neurofilament light (NfL) ELISA was performed using the NF-light® kit from Uman Diagnostics (Quanterix, 10-7001), following the manufacturer’s instructions. The concentrations of NfL were calculated by plotting the average optical density (λ 450 – λ reference) against the standard curve concentrations.
- Immunofluorescence staining of assembloid sections Assembloids were fixed in 4% formaldehyde (Sigma-Aldrich, 1004965000) overnight at 4°C. They were then embedded in 3% low-melting point agarose (Thermo Scientific, R0801) and sectioned into 60μm sections using a Vibratome (Leica, VT1000 S). For immunofluorescence staining, assembloid sections were incubated for 1h in blocking and permeabilization solution (0.5% Triton (Sigma-Aldrich, 93443), 5% goat serum (Invitrogen, 10000C) in 1x PBS) at room temperature. After that, they were incubated for 48h with antibodies (Table S1) in primary antibody solution (0.2% Triton, 5% goat serum, 0.1% Na-Azide (Morphisto, 18189.00100) in 1x PBS) at 4°C. Sections were washed three times for 20 minutes in 0.1% Tween (Sigma-Aldrich, P7949) in PBS at room temperature. Then, they were incubated for 1h with secondary antibodies (Table S1) in secondary antibody solution (0.1% Tween, 5% goat serum, 0.1% Na-Azide in 1x PBS) at room temperature protected from light. Three washes in 0.1% Tween in PBS were done, and sections were rinsed with milliQ H2O and mounted with Fluoromount-G (Thermo Scientific, 00-4958-020) mounting medium. After drying, sections were imaged using a Yokogawa CellVoyager CV8000 High-Content Screening System.

## Results and Discussion

### Patient-derived midbrain assembloids from 3x SNCA patients show increased neuronal death

Midbrain organoids were generated from human iPSCs, as described previously [12], [13]. A wild-type and 3x SNCA iPSC lines (Table S2) were used for the generation of macrophage and neural precursor cells. The latter were used for the generation of MOs, which were then co-cultured with their line-matched macrophage precursor cells for assembloid generation (Figure 1A).

**Figure 1.**
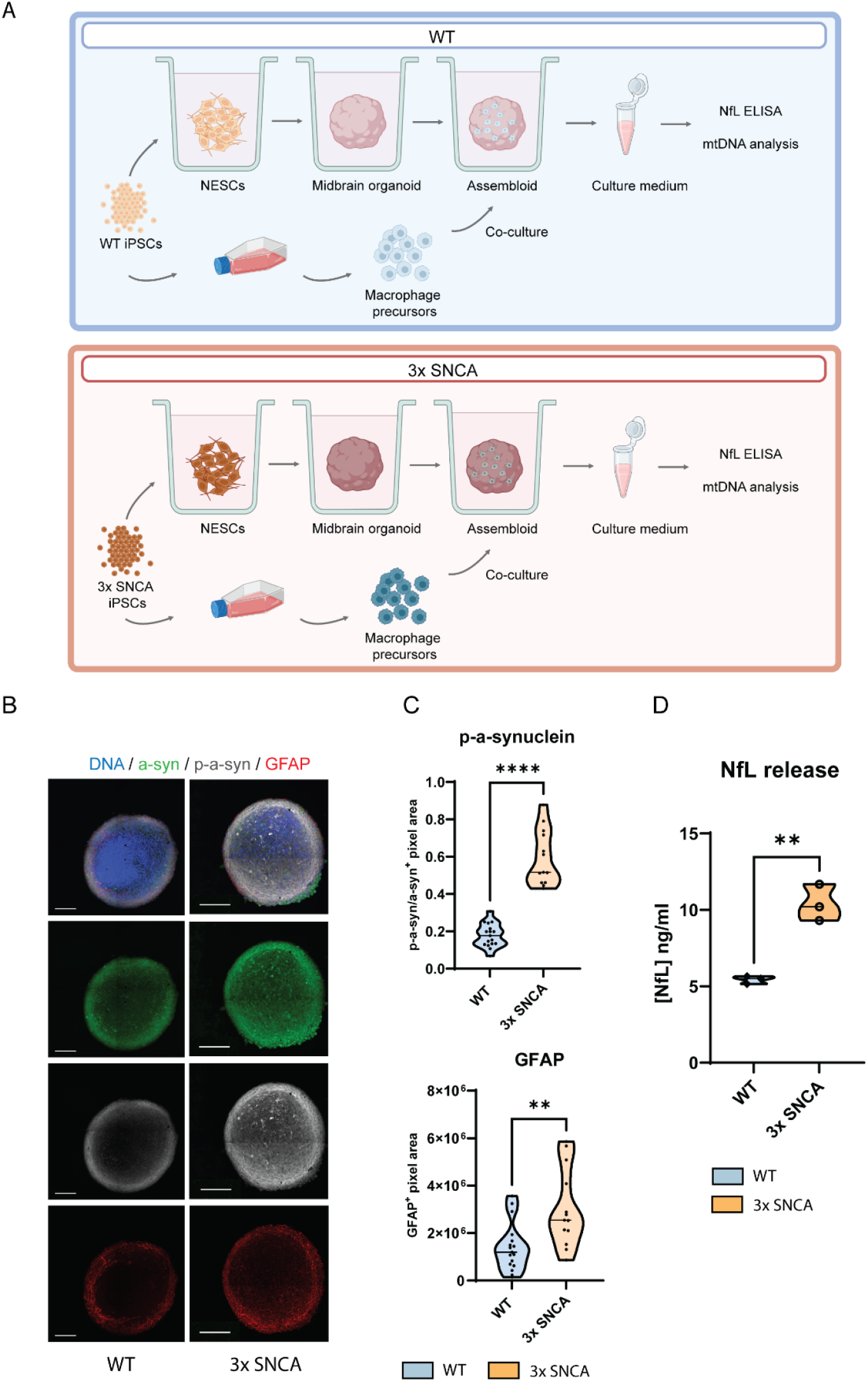
Human midbrain assembloids from the 3x SNCA group show neurodegeneration and synucleinopathy-related phenotypes. A. Diagram showing the culture and analysis methods for the WT group (upper panel) and 3x SNCA (bottom panel). B. Immunofluorescence images showing WT (left panels) and 3x SNCA (right panels) sections stained for alpha-synuclein (a-syn, green), phosphorylated alpha-synuclein (p-a-syn, grey), and GFAP (red). Scale bar = 200μm. C. Violin plots showing the image analysis quantification results for phosphorylated alpha-synuclein (p-a-synuclein, upper graph) and GFAP (bottom graph). Each dot represents an organoid; n = 13-17. D. Violin plot showing released NfL in culture supernatants, measured by ELISA. Each dot represents a cell culture batch; n = 3. ** = p ≤ 0.01; **** = p ≤ 0.0001.

To validate PD-related phenotypes, we performed an immunofluorescence staining of assembloid sections. We used a-syn, p-a-syn, and GFAP antibodies (Fig 1B) and used a published image analysis pipeline to quantify the positive marker area [15]. We observed significantly higher levels of the phosphorylated a-syn proportion to total synuclein in the 3x SNCA assembloids. This indicates a possible alpha-synuclein aggregation in the model, characteristic of PD and other synucleinopathies [16], [17]. Furthermore, the GFAP^+^ area was significantly higher in the 3xSNCA group (Fig 1C). This increased GFAP^+^ area correlates with previous studies on 3x SNCA organoids [18], and could also indicate an astrogliosis process in the PD group.

Assembloids were cultured for 60 days, culture supernatant was collected for neurofilament-light (NfL) release analysis. We observed a significant increase in NfL release in the 3x SNCA group, indicating neurodegeneration (Figure 1D). NfL release has been established as a neurodegeneration and phenotype severity in CSF samples from PD patients [19]This parameter has been detected in cell culture supernatant samples from human cortical organoids [20], however, it had never been observed in patient-specific midbrain organoid or assembloid cultures. For the first time, we report a significant increase in released NfL in midbrain assembloids, demonstrating the clinical translatability of the model, and opening a door to PD biomarker detection in iPSC-derived midbrain assembloids.

### Patient-derived assembloids release higher levels of mitochondrial DNA

Assembloid media was collected for mitochondrial DNA extraction and digital PCR (dPCR) analysis. Three parameters were assessed: mitochondrial DNA copy number, 7S DNA-mediated transcription, and the occurrence of major arc deletions. Via digital PCR, the gene abundance of *ND1* (NADH dehydrogenase 1), a mitochondrial gene, and *B2M* (Beta-2 microglobulin), a nuclear gene, were assessed. The mitochondrial DNA copy number was calculated by determining the ND1/B2M ratio. The copy number levels detected in culture supernatants from the 3x SNCA group were statistically higher than the ones from the WT (Fig. 2A). When normalizing the *ND1* abundance to the assembloid size, we observed a tendency to an increased copy number in the 3x SNCA group (Fig. 2B), following the same trend as the *ND1/B2M* ratio. Since the data normalization to B2M in the medium takes into account a DNA release coming, most likely, from cell death, the *ND1/B2M* ratio gives valuable information on the mitochondrial DNA abundance due to active release rather than cell death. The fact that the differences in *ND1/B2M* are significant and the *ND1/*size are not could highlight a process of active mitochondrial DNA release in the 3x SNCA group. Overall, the findings on the copy number in the media correlate with previous studies that report alterations in mitochondrial homeostasis in PD [21], [22], [23]. Increased blood levels of mitochondrial DNA have been detected in PD patients [21]. Furthermore, higher levels of mitochondrial DNA have been found in genetic PD patients compared to idiopathic forms, leading to a more accentuated neuroinflammatory phenotype [22].

**Figure 2.**
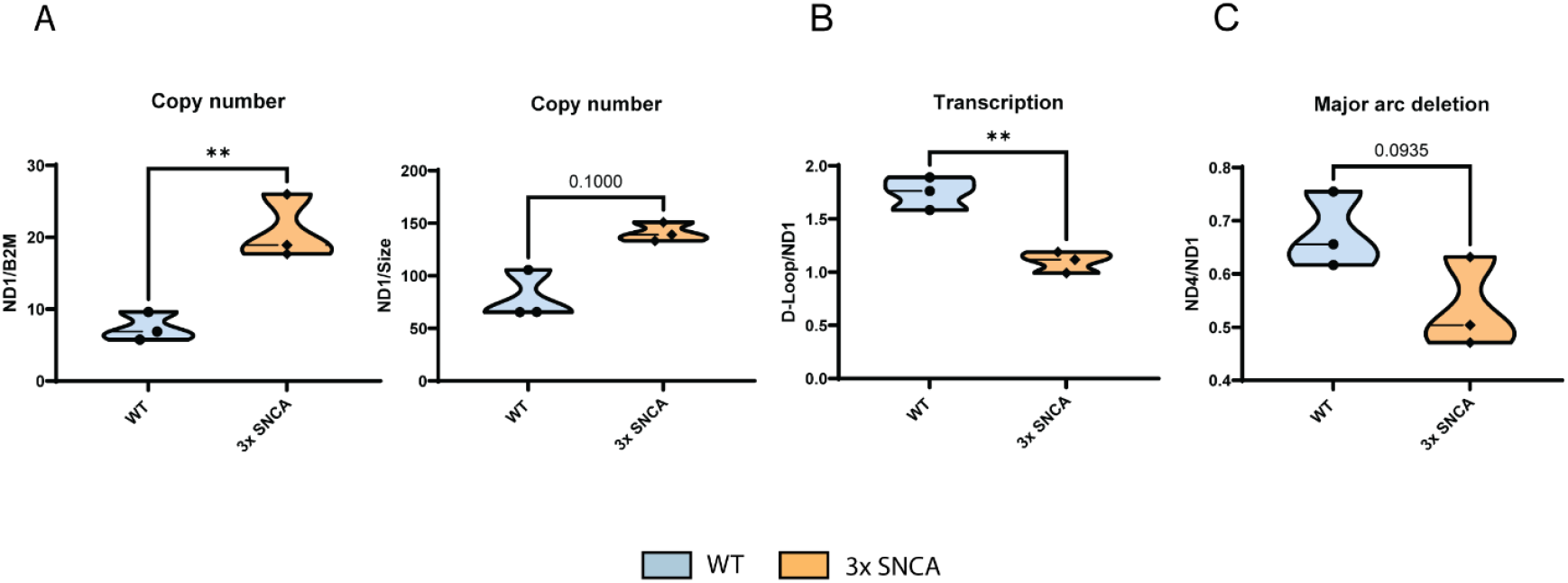
Assembloids from the 3x SNCA group show alterations in mitochondrial DNA dynamics. A. Copy number measured by *ND1/B2M* ratio, showing a significant increase in 3x SNCA supernatant. B. Copy number measured by *ND1*/assembloid size, showing a tendency to an increased ratio in the 3x SNCA group. C. Transcription rate measured by D-Loop*/ND1*, showing lower levels in 3x SNCA assembloid supernatant. D. Major arc deletion ratio measured by *ND4/ND1*, showing a tendency to a decrease in 3x SNCA supernatant. Each dot represents a cell culture batch; n = 3. ** = p ≤ 0.01; ns = non-significant.

Mitochondrial transcription was assessed quantifying the abundance of 7S DNA in the D-loop (displacement loop) region relative to *ND1*. 7S DNA is a fragment, which serves as a natural “primer” during mtDNA transcription ([24]). When present, the D-loop assumes a triple-strand state that can be detected by dPCR. Our results showed a significantly lower transcription rate in the supernatant from the 3x *SNCA* group compared to the WT (Fig. 2C). Finally, the major arc deletion rate was assessed by dividing *ND4* (*NADH dehydrogenase 4*) by *ND1*. Since the *ND4* gene is located in the major arc of the mitochondrial DNA genome, which is commonly subjected to deletions, the ratio *ND4*/*ND1* indicates the deletion rate in mitochondrial DNA. There was no significant difference for this parameter, although a tendency to decrease was observed in the 3x SNCA group, indicating a possible higher deletion number in the 3x SNCA group (Fig. 2D). Significant decrease in mitochondrial transcription (via TFAM expression) as well as multiple deletions [23], [25] have been previously reported in neuronal cultures from post-mortem human PD tissue.

Overall, this study shows how midbrain assembloids are a successful tool for translating clinical phenotypes into *in vitro* research. Furthermore, the fact that the observed phenotypes were detected in cell culture media, with uncomplicated manipulation and short experimental time, indicates that these parameters are not only translatable but also suitable for high-throughput analysis. Finally, these findings have implications for the potential application of NfL and released mitochondrial DNA as biomarkers.

## Conclusion

Human midbrain assembloids are an effective model for detecting neurodegeneration via NfL abundance in cell culture supernatants. Furthermore, alterations in mitochondrial copy number and transcription were detected in supernatants from a 3x SNCA PD patient, which correlates with studies where human PD blood samples were analyzed. In summary, our results indicate that assembloids could be candidate models for detecting PD biomarkers *in vitro* with straightforward techniques compatible with high-throughput analyses.

## Abbreviations

PD: Parkinson’s Disease
MOs: Midbrain Organoids
iPSC: induced Pluripotent Stem Cells
3D: 3-dimensional
2D: 2-dimensional
3x SNCA: Triplication in the SNCA gene
DNA: Deoxyribonucleic acid
dPCR: Digital polymerase chain reaction
NfL: Neurofilament light chain
a-syn: alpha-synuclein
p-a-syn: phosphorylated alpha-synuclein

## Declarations

JS and JJ are co-founders of OrganoTherapeutics SARL.

## Acknowledgments

This project has received funding from the European Innovation Council (Grant No. 101046920 “iSenseDNA”). A.G. and J.G. were supported by a grant from the Michael J. Fox Foundation that investigates “Common Mitochondrial DNA Deletions as a Peripheral Marker of Parkinson’s Disease”.

