## Supplemental Tables for "Released mitochondrial DNA and neurofilament light chain as Parkinson’s disease phenotypes in patient-specific midbrain assembloids"

*Supplementary Information (SI)*

**Table S1 - Antibodies**

| Antibody/Dye name | Species | Manufacturer | Catalog number | Dilution |
|---|---|---|---|---|
| Hoechst | - | Invitrogen | 62249 | 1:10,000 |
| alpha-synuclein | Mouse | Bio-techne | NBP1- 05194 | 1:500 |
| phospho-alpha-synuclein | Rabbit | Abcam | ab51253 | 1:500 |
| GFAP | Chicken | Sigma-Aldrich | AB5541 | 1:1000 |
| Anti-Mouse IgG (H+L), Alexa Fluor 488 | Donkey | Invitrogen | A21202 | 1:1000 |
| Anti-Chicken IgG (H+L), Alexa Fluor 647 | Donkey | Invitrogen | A78952 | 1:1000 |
| Anti-Rabbit IgG (H+L), Alexa Fluor 568 | Donkey | Invitrogen | A10042 | 1:1000 |

**Table S2 - Cell lines**

| Cell line acronym | Disease | Gene | Alteration | Sex | Age of sampling | Source |
|---|---|---|---|---|---|---|
| WT | Healthy | - | - | F | 53 | Reinhardt et al. 2013 |
| 3x SNCA | PD | SNCA | 3x | F | 55 | NINDS |
